## Supplemental Figures for "*C. elegans* Dicer acts with the RIG-I-like helicase DRH-1 and RDE-4 to cleave dsRNA"

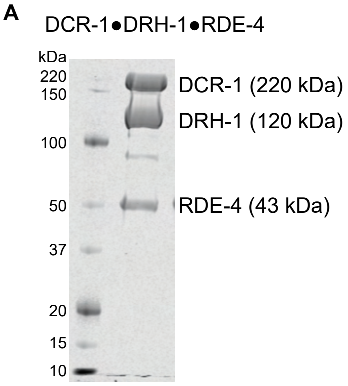


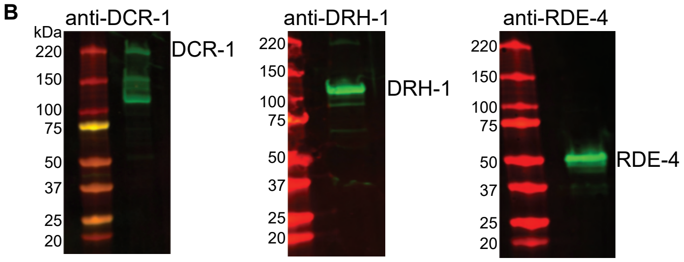


C

| Protein | Amino acid number | Protein Coverage (%) | Number of unique peptides |
|---|---|---|---|
| DCR-1 | 1910 | 77 | 292 |
| DRH-1 | 1091 | 83 | 240 |
| RDE-4 | 385 | 72 | 71 |

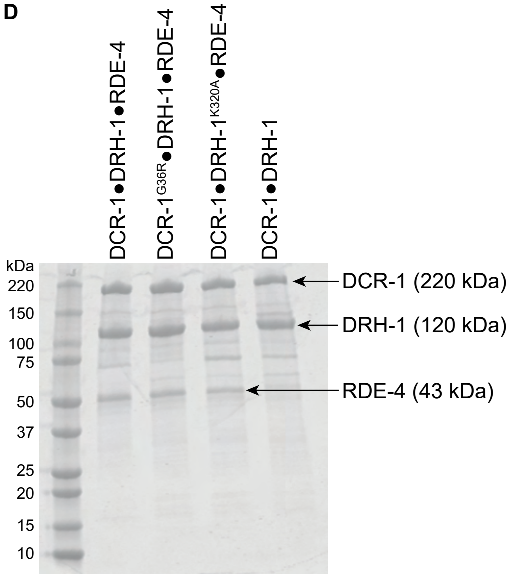


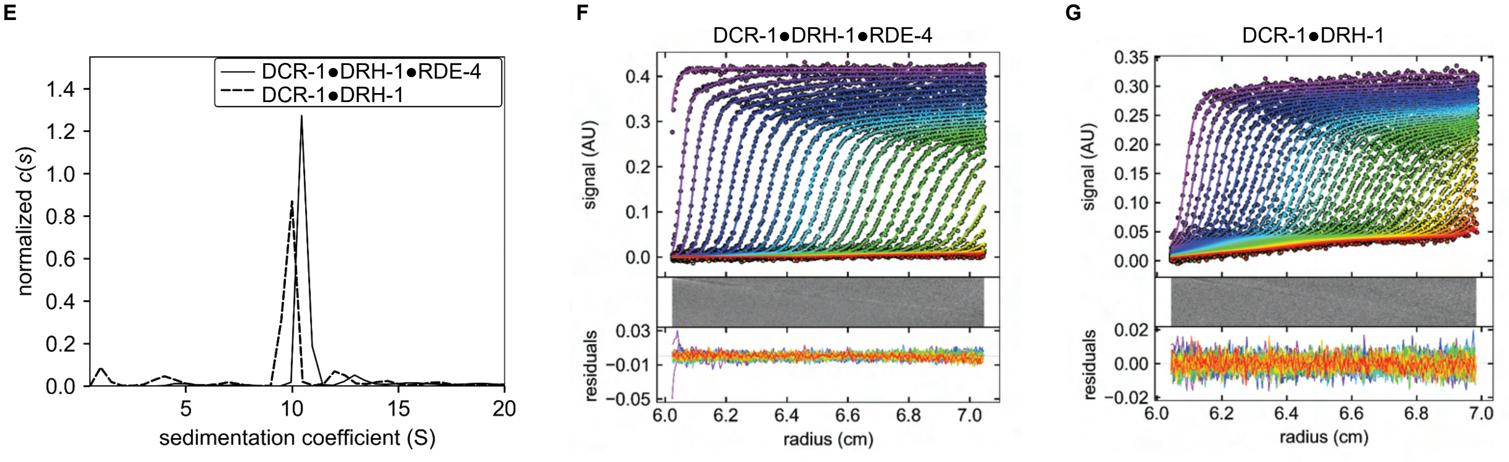


**Figure S1.** Analysis and validation of *C. elegans* antiviral complex after gel filtration. Related to Figure 1.

A. Coomassie-stained SDS-PAGE of DCR-1•DRH-1•RDE-4. Marker molecular weights (kDa) to left of gel and relevant protein names to right.

B. Western blot analyses validating the presence of DCR-1, DRH-1, and RDE-4. Some preparations of Dicer showed lower molecular weight fragments; comparisons to other analyses suggested the commercial antibody preparation detected non-DCR-1 bands, possibly because it was raised against an impure preparation.

C. Mass spectrometry analyses indicating the presence of DCR-1, DRH-1, and RDE-4.

D. Coomassie-stained SDS-PAGE of indicated protein complex. Marker molecular weights (kDa) indicated left of gel and relevant proteins to right.

E-G: SV-AUC sedimentation coefficient (c(s)) analyses of DCR-1•DRH-1•RDE-4 and DCR-1•DRH-1. For E, DCR-1•DRH-1•RDE-4 and DCR-1•DRH-1 complexes were centrifuged at 40,000 RPM, 20ºC, and sedimentation was monitored at 280 nm. C(s) normalized by area showed a s(20,w) = 10.7, f/f0 = 1.75, and MW = 386,284 for DCR-1•DRH-1•RDE-4 , and s(20,w) = 10.1, f/f0 = 1.70, and MW = 336,479 for DCR-1•DRH-1, consistent with 1:1:1 and 1:1 complexes, respectively. DCR-1•DRH-1•RDE-4 (F) and DCR-1•DRH-1(G) absorbance scans and respective fits and residuals, with every second scan and data point shown for clarity (whereas all data were used to obtain fits).

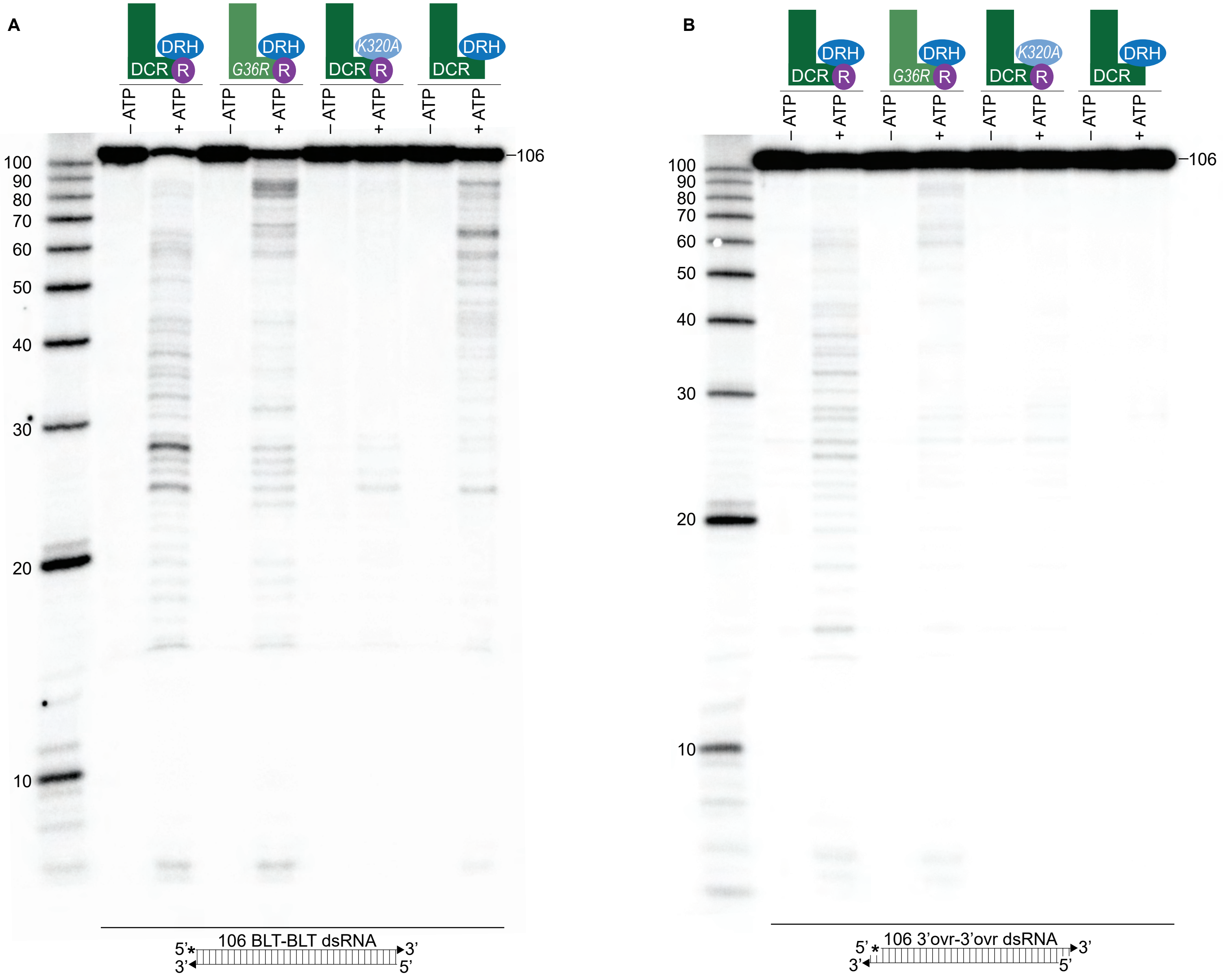
A
DRH
DCR R
G36R R
K320A
DCR
– ATP
+ ATP
100
90
80
70
60
50
40
30
20
10
106
106 BLT-BLT dsRNA
5' 3'
3' 5'
B
106 3'ovr-3'ovr dsRNA

**Figure S2.** 2',3' cyclic phosphates block ATP-independent cleavage and reveal an ATP-dependent side reaction characterized by heterogeneous products. Related to Figure 1.

A. Single-turnover cleavage assays of 106 BLT dsRNA (1nM) with indicated protein complex (50nM) ± 5mM ATP for 60 minutes at 20°C. Sense (top) strand was 5' $^{32}$P-end labeled (*) and both strands contained 2',3' cyclic phosphates (filled triangle in cartoon under gel). Products were separated by 17% denaturing PAGE, and a representative PhosphorImage is shown (n = 3). Left, marker nucleotide lengths. For dmDcr-2, a 2',3' cyclic phosphate at a terminus blocks cleavage from that end[14]. For the antiviral complex, this modification eliminated ATP-independent cleavage (see -ATP lanes), but revealed an ATP-dependent side reaction characterized by a heterogeneous cleavage pattern (see +ATP lanes). Such heterogeneous products required higher concentrations of the antiviral complex, and by optimizing conditions we were able to minimize reaction from the 2',3' cyclic phosphate allowing us to focus on the first cleavage event from the radiolabeled terminus (e.g., see Figures 1C and 1D).

B. Same as A except with 106 3'ovr dsRNA (1nM).

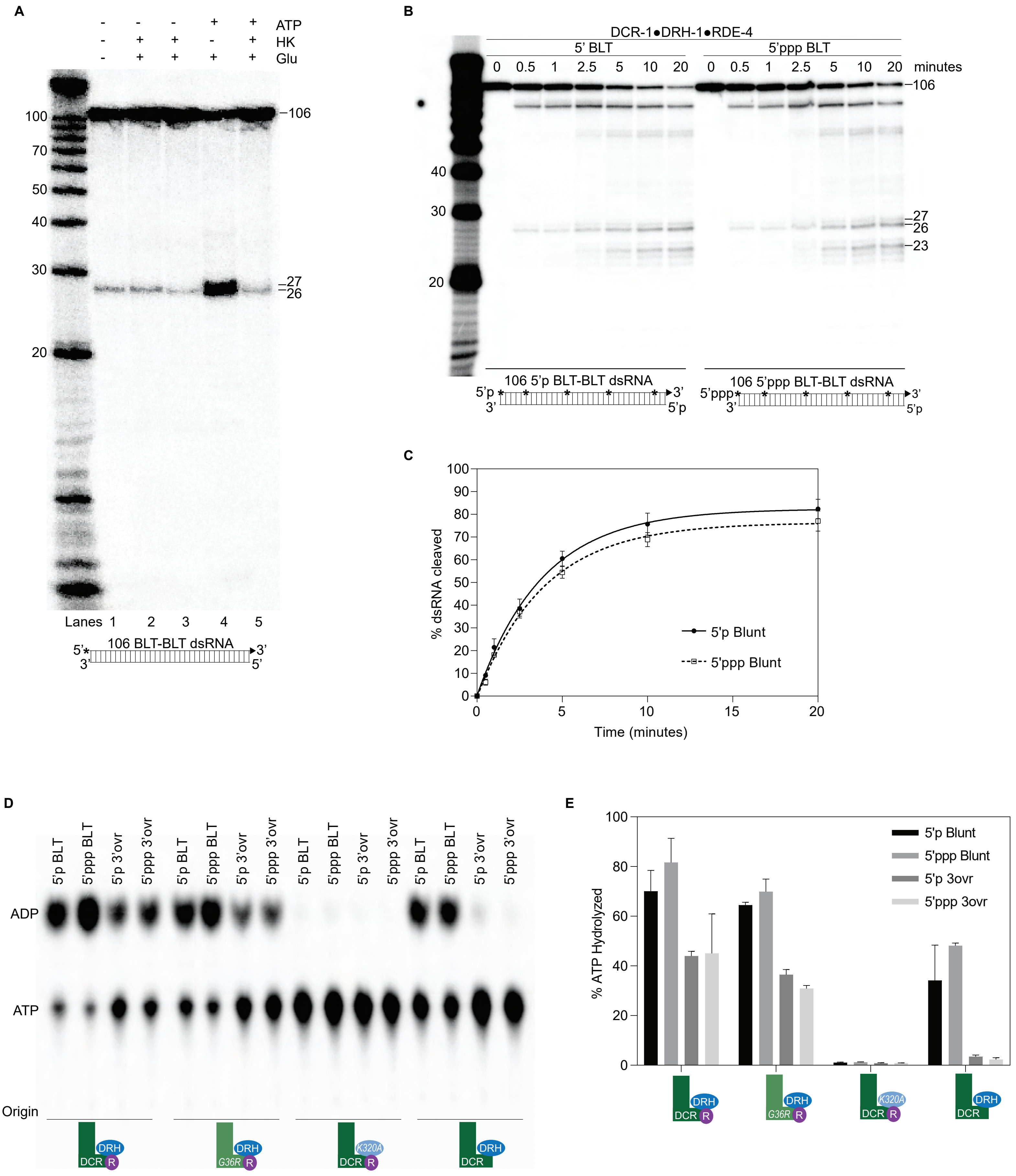
A
ATP
HK
Glu
- - - + +
- + + - +
- + + + +
100
70
50
40
30
20
–106
–27
–26
Lanes 1 2 3 4 5
106 BLT-BLT dsRNA
5'*
3'
3'
5'
B
DCR-1●DRH-1●RDE-4
5' BLT
5'ppp BLT
0 0.5 1 2.5 5 10 20 0 0.5 1 2.5 5 10 20 minutes
–106
40
30
20
–27
–26
–23
106 5'p BLT-BLT dsRNA
5'p
3'
3'
5'p
106 5'ppp BLT-BLT dsRNA
5'ppp
3'
3'
5'p
C
% dsRNA cleaved
Time (minutes)
0 10 20 30 40 50 60 70 80 90 100
0 5 10 15 20
5'p Blunt
5'ppp Blunt
D
5'p BLT
5'ppp BLT
5'p 3'ovr
5'ppp 3'ovr
ADP
ATP
Origin
DRH
DCR
R
G36R
K320A
E
% ATP Hydrolyzed
0 20 40 60 80 100
5'p Blunt
5'ppp Blunt
5'p 3ovr
5'ppp 3ovr
DRH
DCR
R
G36R
K320A

**Figure S3.** DCR-1•DRH-1•RDE-4 cleaves blunt dsRNA in an ATP-independent manner. Triphosphates do not affect cleavage or ATP hydrolysis by DCR-1•DRH-1•RDE-4. Related to Figure 1.

A. Single-turnover cleavage assays of DCR-1•DRH-1•RDE-4 (50nM) with 106 BLT dsRNA (1nM) and ± 5mM ATP, 10mM glucose (Glu), or hexokinase (HK) as indicated for 60 minutes at 20°C. Lane 2 contained 1 unit of HK and lanes 3 and 5 contained 2 units of HK. Sense (top) strand was 5' $^{32}$P-end labeled (*) and contained a 2',3' cyclic phosphate (filled triangle in cartoon under gel). Products were separated by 17% denaturing PAGE, and a representative PhosphorImage is shown (n = 3). Left, marker nucleotide lengths.

B. Single-turnover cleavage assays of DCR-1•DRH-1•RDE-4 (25nM) with 5'p or 5'ppp 106 BLT dsRNA (1nM) at 20°C for times indicated. Sense strand was internally-labeled by including $^{32}$P-ATP during in vitro transcription (*), allowing all cleavage intermediates to be observed. Products were separated by 17% denaturing PAGE, and a representative PhosphorImage is shown. Left, marker nucleotide lengths.

C. Quantification of single-turnover assays as in B. Data points are mean ± SD (n = 3).

D. 50nM of indicated protein complex was incubated with 200nM of 5'p 106 BLT, 5'ppp 106 BLT, 5'p 106 3'ovr, or 5'ppp 106 3'ovr dsRNA with 100μM α-$^{32}$P-ATP at 20°C for 15 minutes. ATP hydrolysis monitored by TLC, and a representative PhosphorImage is shown. Positions of origin, ATP, and ADP are indicated.

E.  Quantification of ATP hydrolysis assays as in D. Data points are mean ± SD (n = 2).

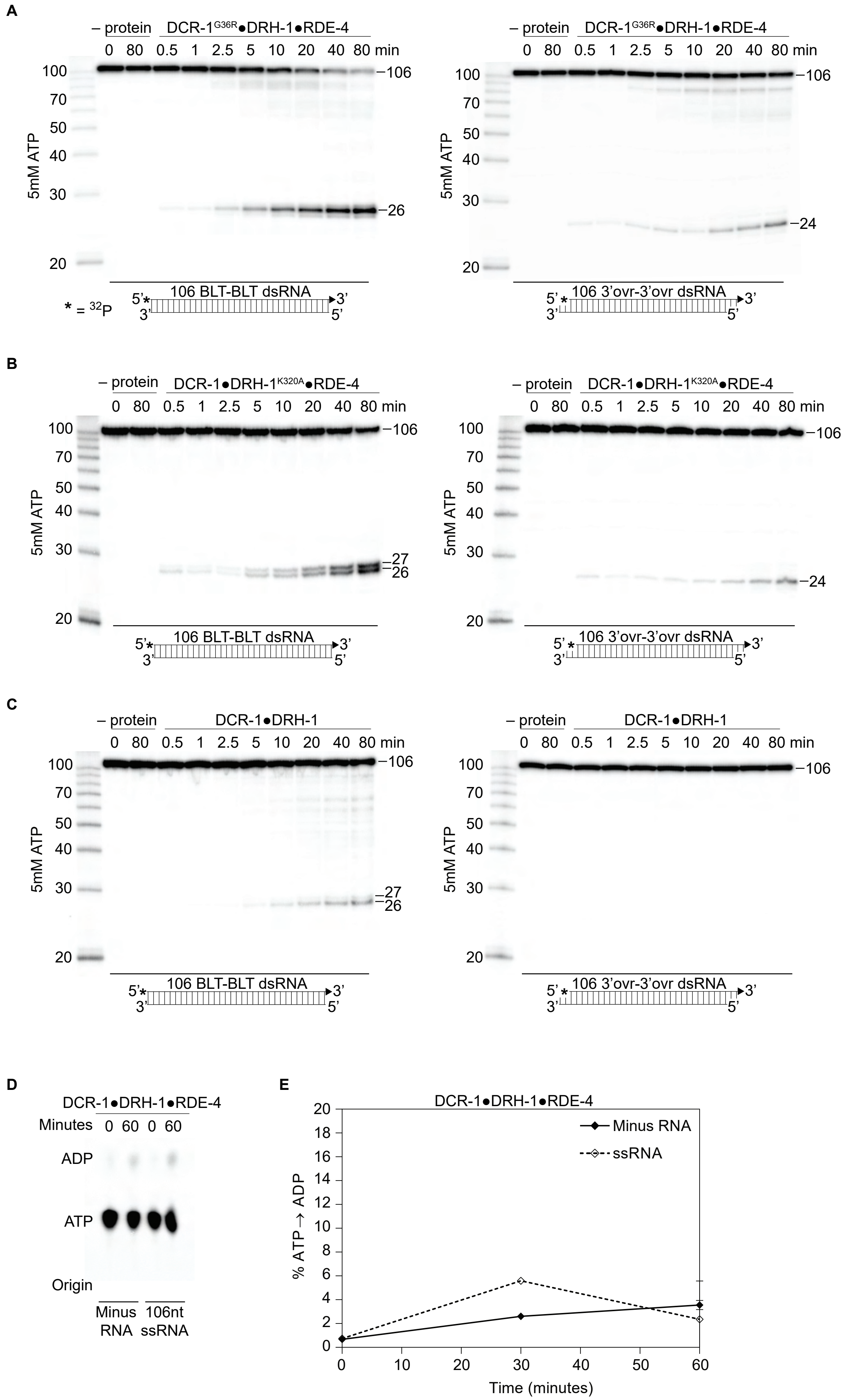

A
– protein
DCR-1G36R●DRH-1●RDE-4
0 80 0.5 1 2.5 5 10 20 40 80 min
5mM ATP
100 70 50 40 30 20
–106
–26
–24
106 BLT-BLT dsRNA
106 3'ovr-3'ovr dsRNA
5' 3' 3' 5'
* = 32P
B
DCR-1●DRH-1K320A●RDE-4
–27
–26
–24
C
DCR-1●DRH-1
D
DCR-1●DRH-1●RDE-4
Minutes 0 60 0 60
ADP
ATP
Origin
Minus RNA
106nt ssRNA
E
DCR-1●DRH-1●RDE-4
Minus RNA
ssRNA
% ATP → ADP
0 2 4 6 8 10 12 14 16 18 20
0 10 20 30 40 50 60
Time (minutes)

**Figure S4.** DCR-1, DRH-1, and RDE-4 affect ATP-dependent cleavage rates. There is a negligible amount of ATP hydrolysis in the absence of dsRNA or presence of ssRNA. Related to Figure 2.

A – C. PhosphorImages show representative primary data (n=3) used for graph shown in Figure 2C. Single-turnover cleavage assays of 25nM DCR-$1^{G36R}$•DRH-1•RDE-4 (A), DCR-1•DRH-$1^{K320A}$•RDE-4 (B) or DCR-1•DRH-1 (C), with 106 BLT or 3'ovr dsRNA (1nM) and 5mM ATP at 20°C. Sense strand was 5' $^{32}$P-end labeled (*). Products were separated by 17% denaturing PAGE. Left, marker nucleotide lengths.

D. DCR-1•DRH-1•RDE-4 (100nM) was incubated without RNA or with 600nM ssRNA and with 100μM α-$^{32}$P-ATP at 20°C. ATP hydrolysis monitored by TLC, and a representative PhosphorImage is shown. Positions of origin, ATP, and ADP are indicated.

E. Quantification of ATP hydrolysis assays as in D. Data points are mean ± SD (n = 3).

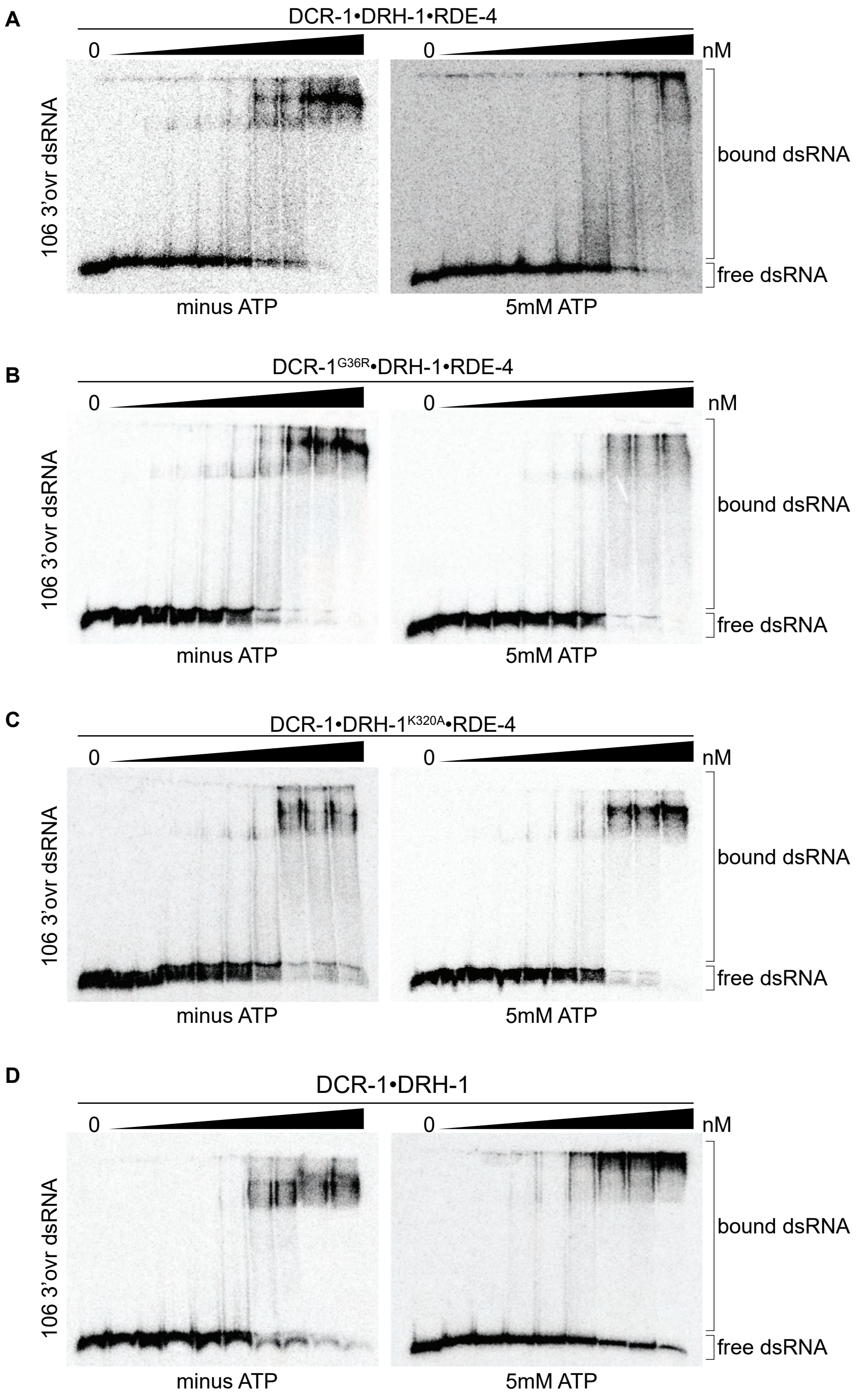

A
DCR-1•DRH-1•RDE-4
0
0
nM
106 3'ovr dsRNA
bound dsRNA
free dsRNA
minus ATP
5mM ATP
B
DCR-1G36R•DRH-1•RDE-4
0
0
nM
106 3'ovr dsRNA
bound dsRNA
free dsRNA
minus ATP
5mM ATP
C
DCR-1•DRH-1K320A•RDE-4
0
0
nM
106 3'ovr dsRNA
bound dsRNA
free dsRNA
minus ATP
5mM ATP
D
DCR-1•DRH-1
0
0
nM
106 3'ovr dsRNA
bound dsRNA
free dsRNA
minus ATP
5mM ATP

**Figure S5.** Binding affinity of DCR-1•DRH-1•RDE-4 wildtype and mutant complexes for 106 3'ovr dsRNA in the absence or presence of ATP. Related to Figure 3.

A. Representative PhosphorImages showing gel mobility shift assays of increasing concentrations of DCR-1•DRH-1•RDE-4 ranging from 0 to 10nM with 106 3'ovr dsRNA ± 5mM ATP as indicated. Sense strand was 5' $^{32}$P-end labeled (*) and contained a 2',3' cyclic phosphate as illustrated in Figure 1C. As labeled on right, all dsRNA that migrated through the gel more slowly than $dsRNA_{free}$ was considered bound.

B – D. Same as A except with indicated complexes. Protein concentrations increased from left to right as indicated and range from 0 to 10nM (B and C), and 0 to 100nM (D).

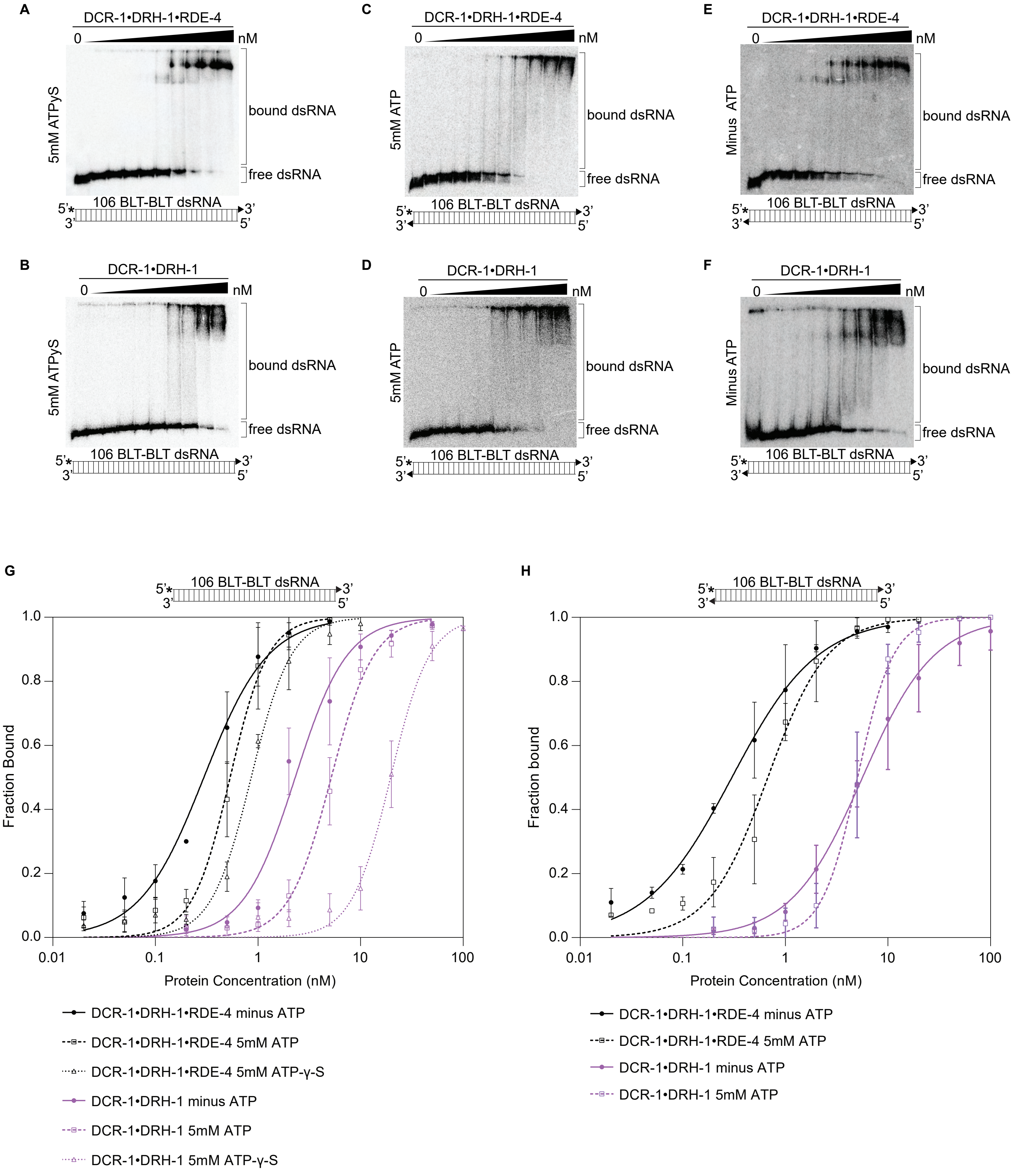
A
DCR-1•DRH-1•RDE-4
0
nM
5mM ATPγS
bound dsRNA
free dsRNA
106 BLT-BLT dsRNA
5'
3'
3'
5'
B
DCR-1•DRH-1
0
nM
5mM ATPγS
bound dsRNA
free dsRNA
106 BLT-BLT dsRNA
C
DCR-1•DRH-1•RDE-4
0
nM
5mM ATP
bound dsRNA
free dsRNA
106 BLT-BLT dsRNA
D
DCR-1•DRH-1
0
nM
5mM ATP
bound dsRNA
free dsRNA
106 BLT-BLT dsRNA
E
DCR-1•DRH-1•RDE-4
0
nM
Minus ATP
bound dsRNA
free dsRNA
106 BLT-BLT dsRNA
F
DCR-1•DRH-1
0
nM
Minus ATP
bound dsRNA
free dsRNA
106 BLT-BLT dsRNA
G
106 BLT-BLT dsRNA
Fraction Bound
0.0
0.2
0.4
0.6
0.8
1.0
0.01
0.1
1
10
100
Protein Concentration (nM)
DCR-1•DRH-1•RDE-4 minus ATP
DCR-1•DRH-1•RDE-4 5mM ATP
DCR-1•DRH-1•RDE-4 5mM ATP-γ-S
DCR-1•DRH-1 minus ATP
DCR-1•DRH-1 5mM ATP
DCR-1•DRH-1 5mM ATP-γ-S
H
106 BLT-BLT dsRNA
Fraction bound
0.0
0.2
0.4
0.6
0.8
1.0
0.01
0.1
1
10
100
Protein Concentration (nM)
DCR-1•DRH-1•RDE-4 minus ATP
DCR-1•DRH-1•RDE-4 5mM ATP
DCR-1•DRH-1 minus ATP
DCR-1•DRH-1 5mM ATP

**Figure S6.** Binding affinity of DCR-1•DRH-1•RDE-4 and DCR-1•DRH-1 for 106 BLT dsRNA in the presence of ATPγS, and for 106 BLT blocked with cyclic phosphate on both ends in the presence and absence of ATP. Related to Figure S2.

A – F. Representative PhosphorImages showing gel mobility shift assays with indicated complexes and the presence (5mM) or absence of ATP or ATPγS as indicated on the left side of the gel. Protein concentrations increased from left to right as indicated and range from 0 to 10nM (A and E), 0 to 20 nM (C), and 0 to 100nM (B, D, and F). As illustrated in the cartoon below each gel, all reactions contained 106 BLT dsRNA with the sense (top) strand 5' $^{32}$P-end labeled (*) and one or both strands containing a 2',3' cyclic phosphate (filled triangles). As labeled on the right, all dsRNA that migrated through the gel more slowly than $dsRNA_{free}$ was considered bound.

G. Binding isotherms showing trends for DCR-1•DRH-1•RDE-4 and DCR-1•DRH-1 with 106 BLT dsRNA containing a 2',3' cyclic phosphate on the sense strand (filled triangle in cartoon above graph) ± 5mM ATP or 5mM ATP-γ-S. Data points, mean ± SD (n ≥ 3).

H. Binding isotherms showing trends for DCR-1•DRH-1•RDE-4 and DCR-1•DRH-1 with 106 BLT dsRNA containing a 2',3' cyclic phosphate on the sense and antisense strands (filled triangle in cartoon above graph) ± 5mM ATP. Data points, mean ± SD (n ≥ 3).

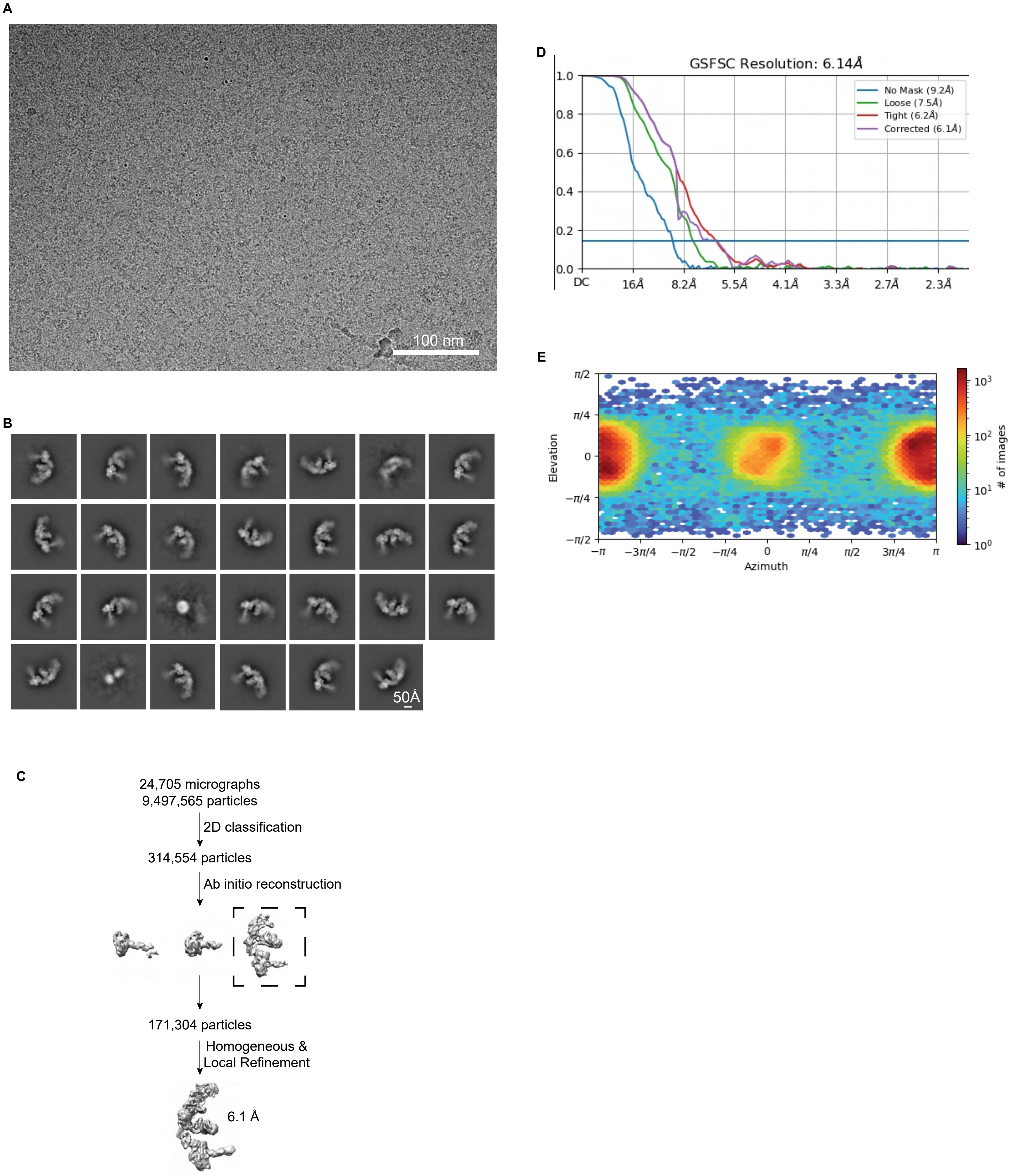
A
100 nm
D
GSFSC Resolution: 6.14Å
No Mask (9.2Å)
Loose (7.5Å)
Tight (6.2Å)
Corrected (6.1Å)
1.0
0.8
0.6
0.4
0.2
0.0
DC
16Å
8.2Å
5.5Å
4.1Å
3.3Å
2.7Å
2.3Å
E
π/2
π/4
0
−π/4
−π/2
Elevation
−π
−3π/4
−π/2
−π/4
0
π/4
π/2
3π/4
π
Azimuth
10^3
10^2
10^1
10^0
### of images
B
50Å
C
24,705 micrographs
9,497,565 particles
2D classification
314,554 particles
Ab initio reconstruction
171,304 particles
Homogeneous &
Local Refinement
6.1 Å

**Figure S7.** Validation of DCR-1•DRH-1•RDE-4 structure with 52 BLT dsRNA, no ATP. Related to Figure 4.

A. Representative cryo-EM micrograph.

B. Representative 2D class averages.

C. Processing tree of cryo-EM data in cryoSPARC.

D. Gold-standard FSC plot.

E. Orientation distribution of particles.

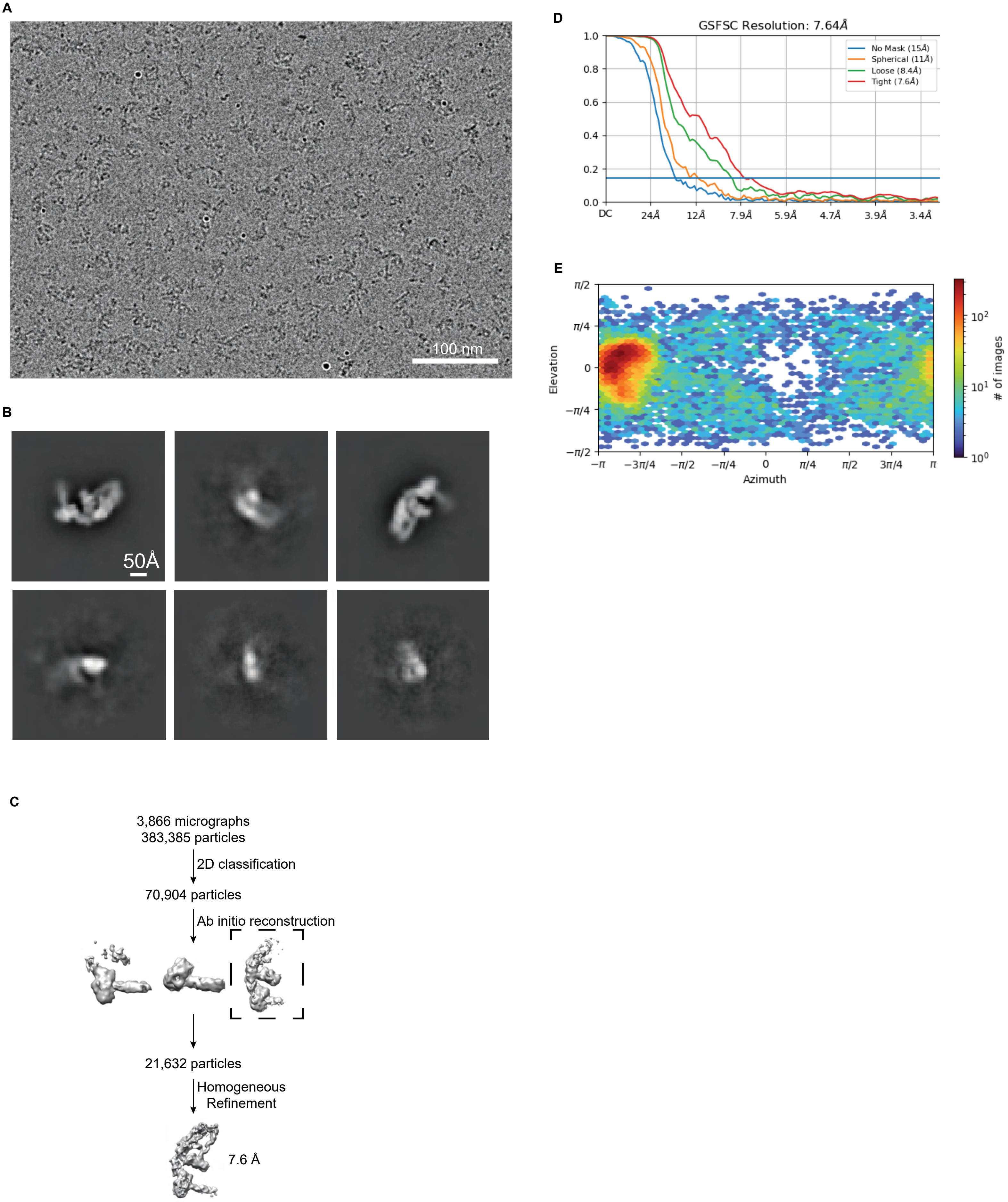
A
100 nm
B
50Å
C
3,866 micrographs
383,385 particles
2D classification
70,904 particles
Ab initio reconstruction
21,632 particles
Homogeneous
Refinement
7.6 Å
D
GSFSC Resolution: 7.64Å
No Mask (15Å)
Spherical (11Å)
Loose (8.4Å)
Tight (7.6Å)
1.0
0.8
0.6
0.4
0.2
0.0
DC
24Å
12Å
7.9Å
5.9Å
4.7Å
3.9Å
3.4Å
E
π/2
π/4
0
−π/4
−π/2
Elevation
−π
−3π/4
−π/2
−π/4
0
π/4
π/2
3π/4
π
Azimuth
10^2
10^1
10^0
### of images

**Figure S8.** Validation of DCR-1•DRH-1•RDE-4 structure with 42 BLT dsRNA. Related to Figure 4.

A. Representative cryo-EM micrograph.

B. Representative 2D class averages.

C. Processing tree of cryo-EM data in cryoSPARC.

D. Gold-standard FSC plot.

E. Orientation distribution of particles.

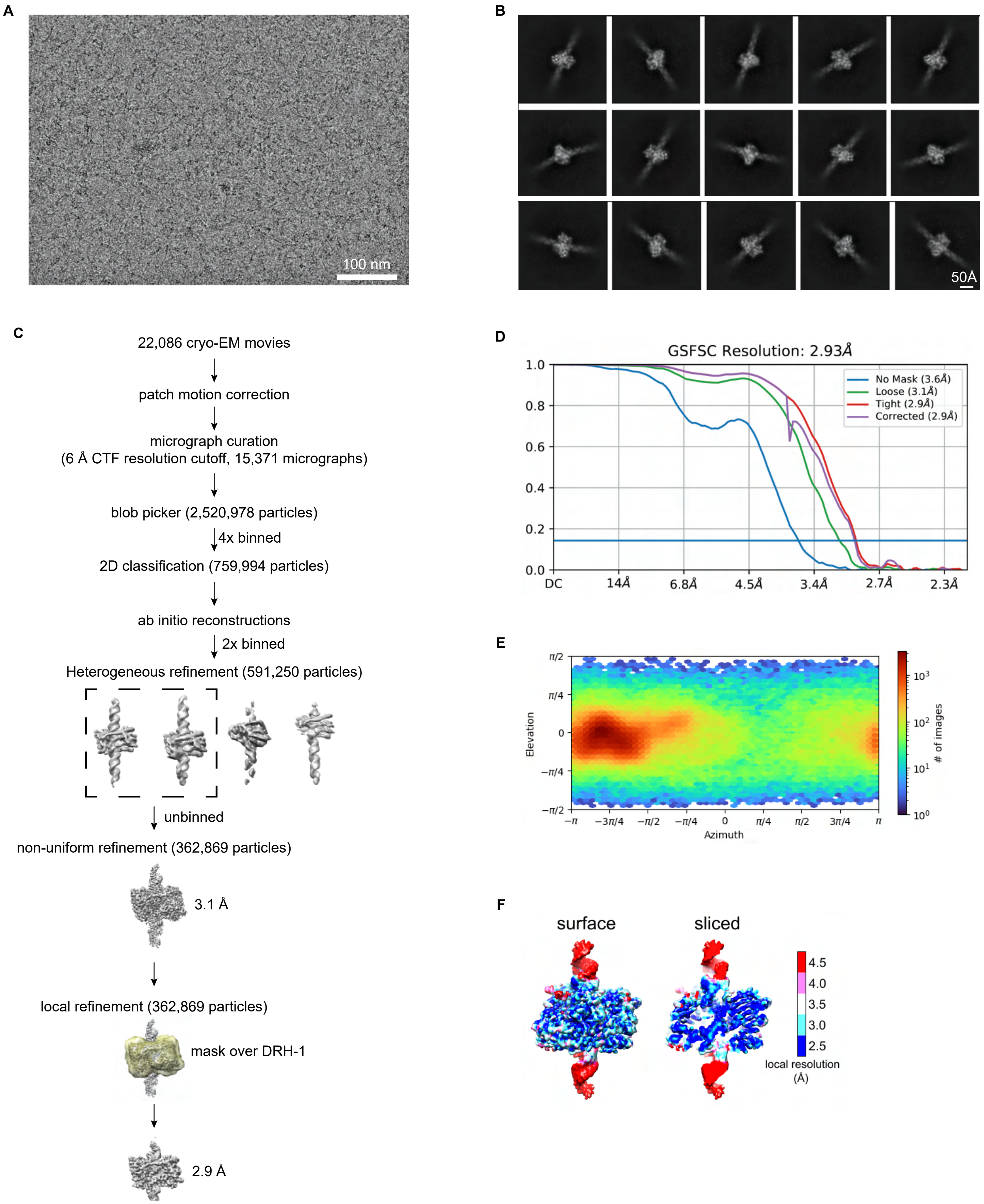
A
100 nm
B
50Å
C
22,086 cryo-EM movies
patch motion correction
micrograph curation
(6 Å CTF resolution cutoff, 15,371 micrographs)
blob picker (2,520,978 particles)
4x binned
2D classification (759,994 particles)
ab initio reconstructions
2x binned
Heterogeneous refinement (591,250 particles)
unbinned
non-uniform refinement (362,869 particles)
3.1 Å
local refinement (362,869 particles)
mask over DRH-1
2.9 Å
D
GSFSC Resolution: 2.93Å
No Mask (3.6Å)
Loose (3.1Å)
Tight (2.9Å)
Corrected (2.9Å)
1.0
0.8
0.6
0.4
0.2
0.0
DC
14Å
6.8Å
4.5Å
3.4Å
2.7Å
2.3Å
E
Elevation
Azimuth
### of images
F
surface
sliced
4.5
4.0
3.5
3.0
2.5
local resolution
(Å)

**Figure S9.** Validation of DRH-1 structure with 106 3'ovr dsRNA. Related to Figure 4.

A. Representative cryo-EM micrograph.

B. Representative 2D class averages.

C. Processing tree of cryo-EM data in cryoSPARC.

D. Gold-standard FSC plot.

E. Orientation distribution of particles.

F. Local resolution heat map.

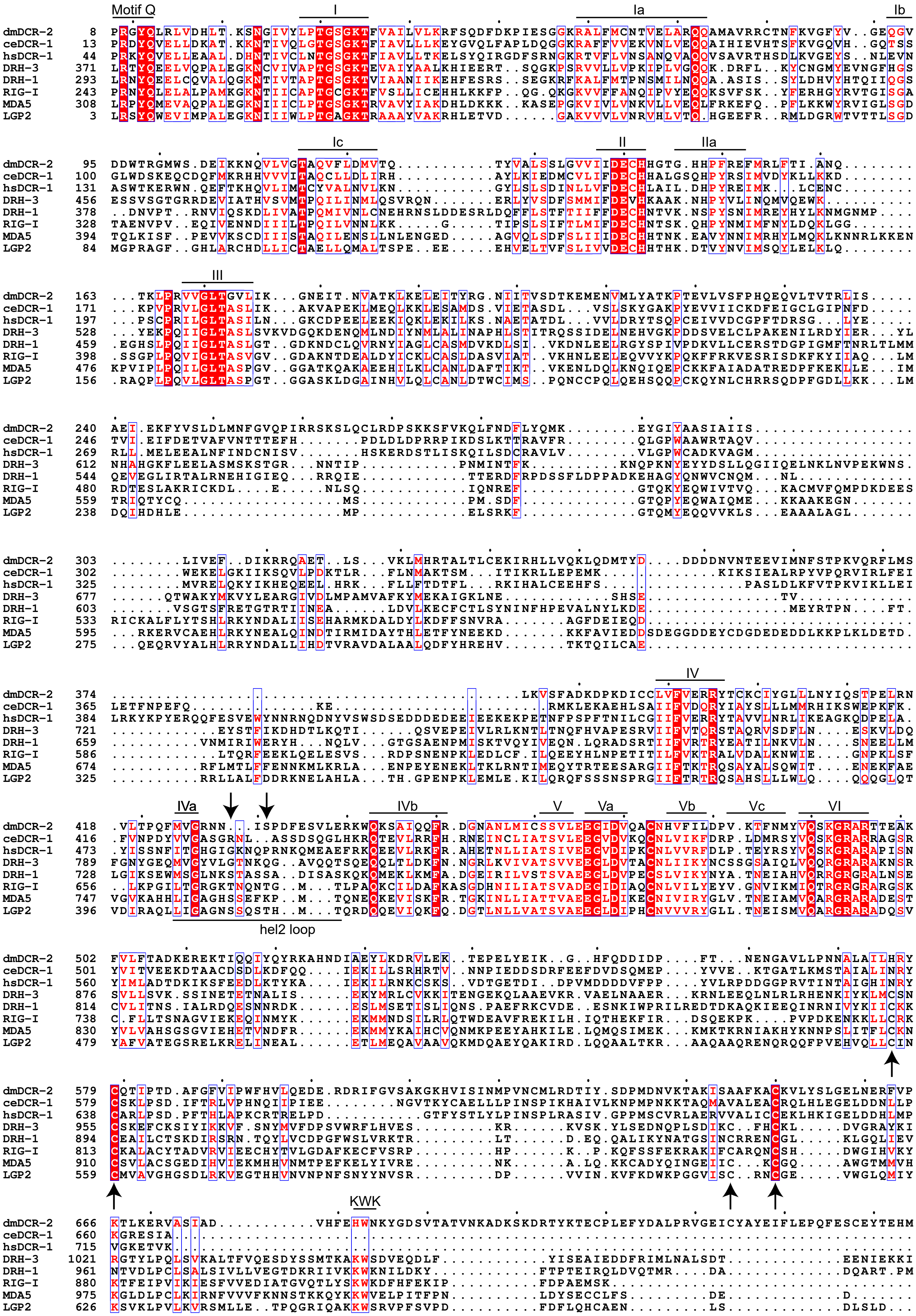

Motif Q
I
Ia
Ib
Ic
II
IIa
III
IV
IVa
IVb
V
Va
Vb
Vc
VI
hel2 loop
KWK
dmDCR-2
ceDCR-1
hsDCR-1
DRH-3
DRH-1
RIG-I
MDA5
LGP2

**Figure S10.** Sequence alignment of RLRs and Dicer helicase and DUF domains. Related to Figure 4.

A. Structure-based multiple sequence alignment was carried out using Clustal Omega Multiple Sequence Alignments[66–68], and illustrated with ESPript[70]. Conserved helicase motifs are indicated with a line above the sequence. Arrows pointing down indicate the two threonine residues phosphorylated in RIG-I. Arrows pointing up indicate conserved cysteines in RLRs.

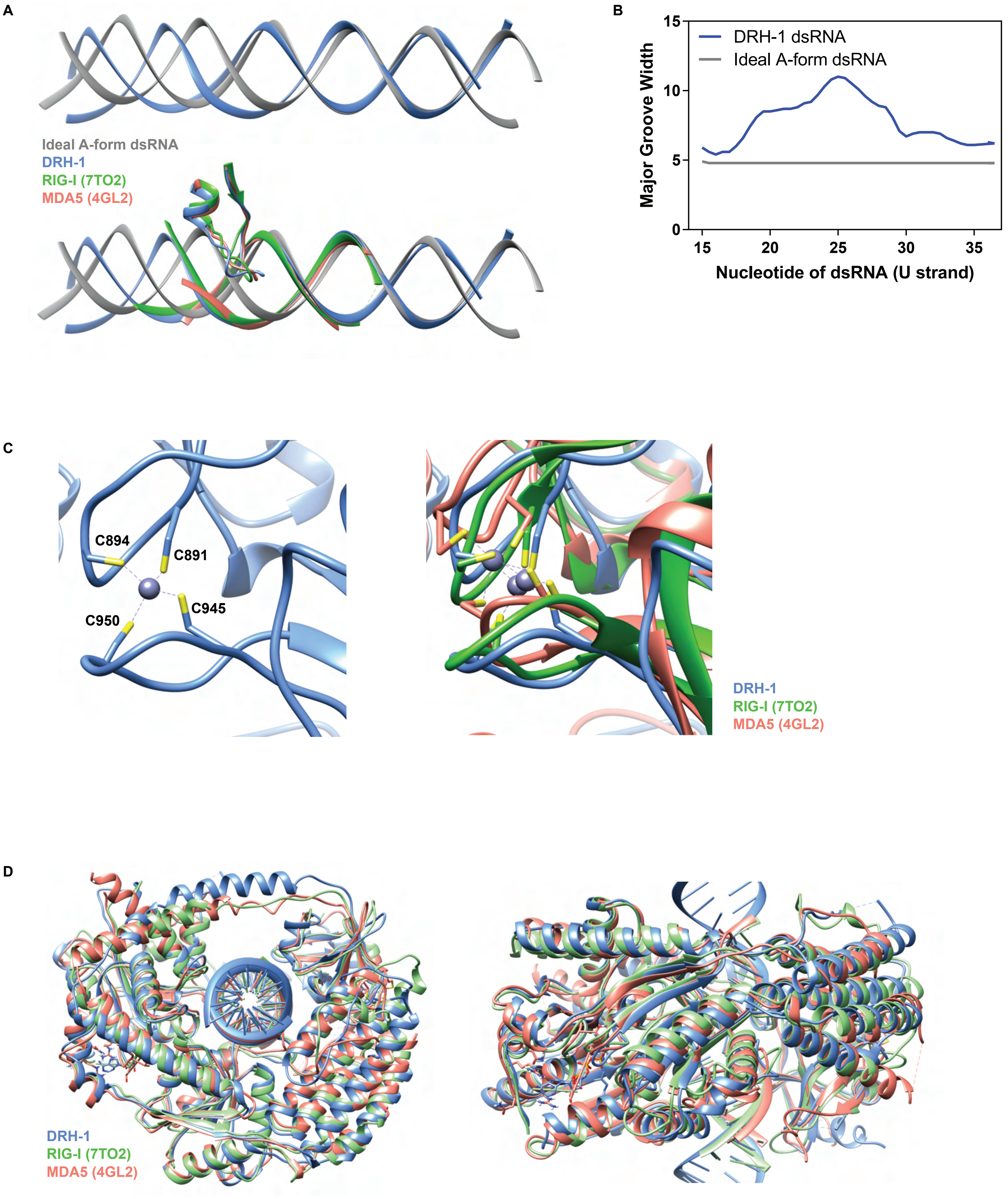
A
Ideal A-form dsRNA
DRH-1
RIG-I (7TO2)
MDA5 (4GL2)
B
DRH-1 dsRNA
Ideal A-form dsRNA
Major Groove Width
15
10
5
0
15
20
25
30
35
Nucleotide of dsRNA (U strand)
C
C894
C891
C945
C950
DRH-1
RIG-I (7TO2)
MDA5 (4GL2)
D
DRH-1
RIG-I (7TO2)
MDA5 (4GL2)

**Figure S11.** Comparison of DRH-1 with RIG-I and MDA5. Related to Figure 4.

A. Top: Comparison between an ideal A-form dsRNA (generated in Chimera) and DRH-1 bound dsRNA. Bottom: Comparison between an ideal A-form dsRNA (generated in Chimera) and dsRNA bound to DRH-1 (blue), RIG-I (green, PDB 7TO2), and MDA5 (salmon, PDB 4GL2).

B. Comparison of the major groove width between an ideal A-form dsRNA and DRH-1 bound dsRNA using Curves+[71,72]. Nucleotides numbered as in our PDB (8T5S).

C. Left: Four cysteines coordinate binding a zinc ion in DRH-1 (blue). Right: Comparison of structural zinc ion between DRH-1 (blue), RIG-I (green, 7TO2), and MDA5 (salmon, 4GL2).

D. Structural alignment of DRH-1 (blue) with RIG-I (green, 7TO2) and MDA5 (salmon, 4GL2).

**Table S1. Summary of $K_d$ values**

| | Kd (nM) | |
|---|---|---|
| | **DCR-1•DRH-1•RDE-4** | **DCR-1•DRH-1** |
| 106 BLT + 5mM ATPγS | 0.86 ± 0.06 | 19.63 ± 1.67 |
| 106 BLT 2'3' cyclic phosphate on both ends minus ATP | 0.30 ± 0.04 | 5.65 ± 0.71 |
| 106 BLT 2'3' cyclic phosphate on both ends + 5mM ATP | 0.67 ± 0.09 | 5.06 ± 0.40 |

Values shown are mean ± SD (n = 3).

| **Table S2. Cryo-EM data collection, refinement, and validation statistics** | | | |
|---|---|---|---|
| **Data Collection and processing** | | | |
| | DRH-1 | Complex bound to one dsRNA | Complex bound to two dsRNA |
| EM Databank Accession ID | EMD-41060 | EMD-43430 | EMD-43431 |
| Microscope | Titan Krios G3 | Titan Krios G3 | Titan Krios G3 |
| Voltage (kV) | 300 | 300 | 300 |
| Detector | Gatan K3 | Gatan K3 | Gatan K3 |
| Data collection software | SerialEM | Leginon | SerialEM |
| Nominal magnification | 81,000x | 81,000x | 81,000x |
| Total number of frames | 40 | 40 | 50 |
| Total electron exposure ($e^-$/Å) | 38 | 50 | 50 |
| Defocus range (µm) | -1.0 - -2.0 µm | -1.0 - -2.0 µm | -1.0 - -2.0 µm |
| Pixel size (Å) | 0.529 Å | 0.533 Å | 0.394 Å |
| Number of micrographs | 14,602 | 24,705 | 3,866 |
| Final particles | 362,869 | 171,304 | 26,879 |
| Resolution (GSFSC 0.143) | 2.9 Å | 6.1 Å | 7.6 Å |
| **Refinement and validation statistics for 2.9 Å reconstruction of DRH-1** | | | |
| **Structure** | | | |
| Protein Data Bank Accession ID | 8T5S | | |
| **Symmetry imposed** | | | |
| 3D classification | C1 | | |
| 3D refinement | C1 | | |
| Final particles | 362,869 | | |
| **Map resolution (Å)** | | | |
| FSC 0.143 (unmasked) | 3.6 Å | | |
| FSC 0.143 (masked, corrected) | 2.9 Å | | |
| **Model Refinement** | | | |
| Initial model used (AlphaFold database) | AF-G5EDI8-F1 | | |
| Map correlation coefficient | 0.8 | | |
| **Model composition** | | | |
| Non-hydrogen atoms | 6779 | | |
| Protein residues | 693 | | |
| Nucleotides | 60 | | |
| Ligands (ADP, $Mg^{2+}$, $Zn^{2+}$) | 3 | | |
| **R.m.s. deviations** | | | |
| Bond lengths (Å) | 0.002 | | |
| Bond angles (°) | 0.499 | | |
| **Validation** | | | |
| MolProbity score | 1.72 | | |
| Clashscore | 10.02 | | |
| Poor rotamers (%) | 0.48 | | |
| **Ramachandran plot** | | | |
| Favored (%) | 96.79 | | |
| Allowed (%) | 3.21 | | |
| Disallowed (%) | 0.00 | | |
| C-beta deviations (0.25 Å) | 0.00 | | |
| CaBLAM outliers (%) | 2.81 | | |
